# Decoupling Fabrication from Encoding: DNA-Addressable Template Microparticles for Large, User-Defined Optical Barcode Libraries

**DOI:** 10.64898/2026.05.10.723434

**Authors:** Akihiro Eguchi, Yuichiro Iwamoto, Hinata Tokuda, Haruka Narita, Adrian M. Martin, Sadao Ota

**Author notes:** These authors contributed equally to this work (Akihiro Eguchi, Yuichiro Iwamoto, Hinata Tokuda, Haruka Narita).

## Abstract

Optical barcodes for pooled high-throughput screening must support large libraries while remaining decodable in a single imaging step. Existing approaches often trade design control for manufacturability: deterministic barcodes often require per-code redesign of particle fabrication, whereas stochastic combinatorial barcodes are difficult to generate as predefined batches. Here we introduce a chemically programmable barcoding architecture that decouples particle fabrication from barcode assignment. Using a contact-free multilaminar flow lithography platform with all-around three-dimensional sheathing, we continuously fabricate a universal hydrogel scaffold containing five spatially segregated DNA-addressable domains at rates >10^6^ particles/h. Chosen barcode identities are subsequently written on demand onto the same template batch by domain-selective DNA hybridization. Single-domain measurements resolved 64 candidate optical states, indicating an experimentally informed theoretical upper bound of 64^5^ ≈ 1.1 × 10^9^ barcodes. We further implemented a predefined 59,049-code library by split-pool labeling, achieving an 88% recovery of decoded beads at a stringent posterior threshold (>0.95). After 11 days, >7,800 beads were correctly re-identified at >0.95 accuracy in matched fields of view. This strategy provides a highly scalable, chemically programmable route to build large, user-defined optical barcode libraries with single-image optical readout and longitudinal traceability.

## Introduction

High-throughput screening increasingly relies on pooled workflows in which individual particles, reaction compartments, or payload carriers must be linked to machine-readable identities that can be decoded rapidly in situ and revisited over time.^[1–5]^ Optical barcodes are attractive for this purpose because they enable nondestructive readout from individual objects with traceability in the imaging field. In practice, however, the number of optical states that can be robustly distinguished from any single barcode feature is limited by spectral crosstalk, signal-intensity variation, and measurement noise. Large optical barcode libraries that remain decodable in a single imaging step therefore require multiple independent barcode elements per particle, rather than assigning an increasing number of closely spaced states to a single optical feature.

Existing multielement barcoding strategies face a persistent trade-off between design-defined barcode identity and scalable library generation. In deterministic particle barcodes, code identity is defined during fabrication through particle geometry, prescribed spatial features, or fixed material composition.^[6–22]^ While this approach provides designed, readily decodable barcodes, expanding the codebook requires redesign of fabrication patterns, flow configurations, or material states for additional codes. Conversely, stochastic approaches expand diversity through random combinatorial incorporation or mixing of barcode elements.^[23–25]^ This approach can efficiently populate large code spaces, but the identity of each particle is set by stochastic assembly rather than being defined by design. Consequently, specific barcode species are difficult to generate on demand in defined quantities, complicating controlled library composition and one-to-one mapping between barcodes and chosen payloads.

What is needed, therefore, is a manufacturable particle architecture that combines the coding efficiency of multielement barcodes with the ability to specify barcode identity by design, as in deterministic single-particle barcodes. We reasoned that this can be achieved if barcode identity is encoded by assigning one of N optical states to each of L ordered, independently addressable domains within a common template particle. This architecture expands coding capacity combinatorially, because assigning one of N states to each of L domains yields N^L^ distinguishable codes within a single-particle format. It also enables direct preparation of defined batches of chosen codes from a common batch of identical templates, thereby decoupling particle fabrication from code assignment. Because the barcode is embedded in fixed domains of a single particle, the same architecture is naturally compatible with spatial fiducials, error-aware decoding, and longitudinal re-identification.

Here we implement this concept using hydrogel template microparticles that contain five spatially segregated DNA-addressable domains (Figure 1). Each domain is defined by a distinct immobilized oligonucleotide handle, allowing barcode states to be written independently after particle fabrication by selective hybridization of fluorophore-labeled complementary strands. Barcode identity is therefore not fixed during fabrication, but assigned post-fabrication across a universal particle scaffold. This architecture combines a common particle template with chemically programmable, domain-resolved encoding.

**Figure 1.**
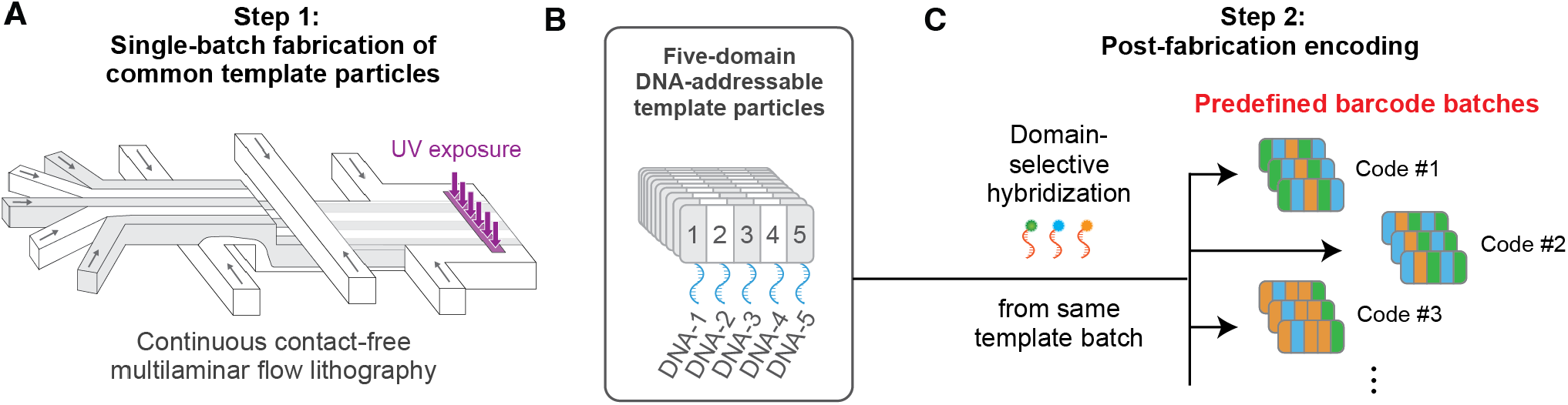
Large-scale programmable optical barcoding by decoupling template-particle fabrication from barcode assignment. (A) Singlebatch, high-throughput fabrication of common hydrogel template particles containing five spatially segregated oligonucleotide domains by continuous contact-free multilaminar flow lithography at rates of >10^6^ particles/h. (B) The common template particles have five distinct layers, each functionalized with an orthogonal DNA sequence. (C) Post-fabrication assignment of barcode identities by domain-selective hybridization of fluorophore-labeled complementary strands, enabling the same template batch to generate predefined barcode sets.

A key challenge, however, is manufacturing such multidomain DNA-addressable templates at scale while preserving domain-level segregation. To address this challenge, we developed a contact-free multilaminar flow lithography platform with all-around three-dimensional sheathing. The platform hydrodynamically isolates the photopolymerization zone from the channel walls and enables continuous fabrication of five-domain particles with spatially segregated DNA-addressable domains at >10^6^ particles h^−1^. We then assigned barcode identities by domain-selective hybridization, including predefined barcode batches from a common template and a 59,049-code (9^5^) library generated by split-pool labeling. Under a stringent posterior decoding threshold (>0.95), 88% of analyzed beads were successfully decoded, and barcode identities remained sufficiently stable for reliable longitudinal re-identification after 11 days. Together, these results establish a practical route to large, user-defined optical barcode libraries that remain decodable in a single imaging step. The same template architecture also provides a chemically addressable foundation for future integration with functional payloads in screening workflows.

## Results and Discussion

### High-throughput Fabrication of Five-domain DNA-addressable Template Microparticles via Contact-free Multilaminar Flow Lithography

The first experimental requirement for the barcoding architecture in Figure 1 is a scalable source of common template particles containing five independently addressable DNA domains. These particles must satisfy two conditions simultaneously: (i) they must preserve domain-level segregation during fabrication so that each DNA-domain can later be addressed independently, and (ii) they must be manufacturable at scale in a continuous, high-throughput process compatible with large barcode libraries.

Flow lithography is a promising route to high-throughput fabrication of spatially patterned functional hydrogel particles.^[26–28]^ In particular, hydrodynamic focusing lithography (HFL) mitigates polymerization near the channel walls and the resulting risk of clogging by hydrodynamically separating the reactive stream from the device walls.^[29]^ To date, multi-domain HFL has predominantly been implemented in a stop-flow configuration at a fabrication rate of 10^3^–10^4^/h.^[17,30]^ However, the overall monomer dwell times inherent to stop-flow operation (typically around 500–1000 ms^[17,28,30]^) permit molecular diffusion across laminar interfaces. For oligonucleotide-functionalized systems, this diffusion can broaden inter-domain boundaries and compromise the domain-level spatial segregation required for reliable post-fabrication optical encoding. Moreover, we are not aware of reports demonstrating continuous HFL with more than three stacked laminar layers under continuous photopolymerization, highlighting the practical challenge of maintaining stable multilaminar streams in this regime.

To address these constraints, we designed a two-layer PDMS device that is simple to fabricate and implements all-around 3D sheathing for a five-stream reactive core (Figure 2A). Bottom, top, and lateral sheath flows progressively surround the multilaminar sample stream and hydrodynamically isolate it from the channel walls before UV exposure (bottom panels (i)-(iv) of Figure 2A). This geometry enables continuous photopolymerization while maintaining the multilaminar organization at the polymerization region.

**Figure 2.**
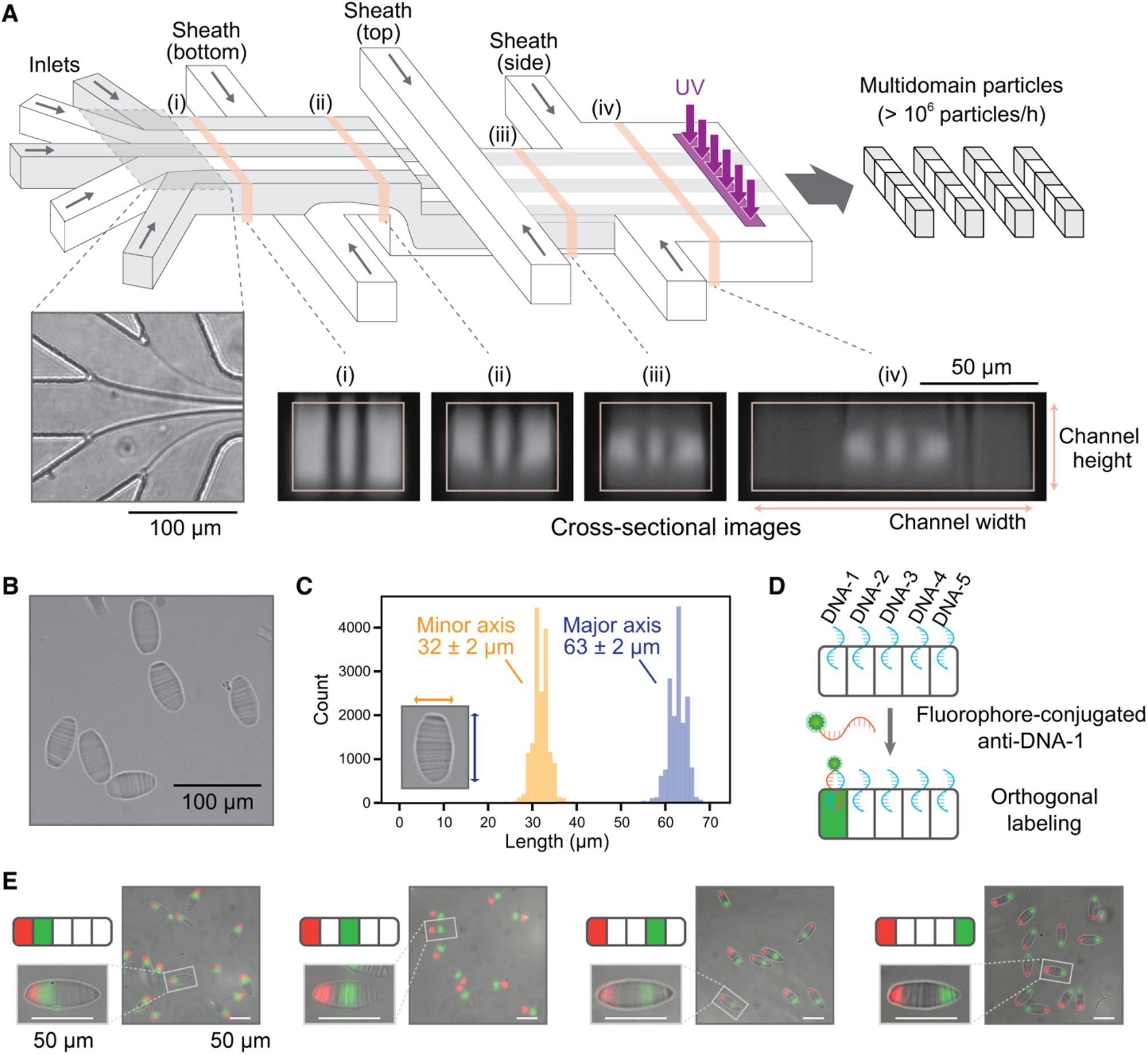
Contact-free multilaminar flow lithography enables scalable fabrication of five-domain DNA-addressable template microparticles. (A) Two-layer PDMS device design. Five reactive inlet streams form a stacked multilaminar core that is progressively surrounded by bottom, top, and lateral sheath flows (i–iv), generating a contact-free reactive stream at the UV polymerization region; inset: bright-field image of the multilaminar junction and representative cross-sectional images along the channel. (B) Representative bright-field image of fabricated fivedomain hydrogel template microparticles. (C) Size distributions of the major and minor particle axes measured from bright-field images. (D) Schematic of domain-specific hybridization of fluorophore-conjugated strands to particles bearing distinct immobilized anchor oligonucleotides. (E) Fluorescence/bright-field overlays showing domain-specific labeling of the intended domains after hybridization with fluorophore-conjugated complementary strands; in each example, domain 1 was labeled with AF647 and one of domains 2–5 was labeled with AF488.

Using this device with spatially patterned strong 500-µs UV pulses, we continuously generated five-domain hydrogel microparticles at rates exceeding 10^6^ particles/h (Figure 2B). Stable operation was maintained for more than 3 hours of continuous fabrication, yielding over 3 million beads in a single run. Bright-field image analysis of the fabricated microparticles showed that the particles were highly uniform, with major and minor axes of 63 ± 2 µm and 32 ± 2 µm, respectively (Figure 2B, 2C), demonstrating that the reactive core stream can be stably maintained during continuous, long-term fabrication. The short residence time before polymerization helps limit inter-domain mixing. Under the flow conditions, the residence time in the lamination region was estimated to be approximately 42 ms (see Methods). Even using an upper-bound diffusion coefficient for single-stranded DNA in aqueous buffer (D ≈ 1.5 × 10^-10^ m^2^/s),^[31]^ the estimated diffusion length over this interval is ~3.5 µm, remaining below the 5 µm separation later used for domain segmentation in the decoding workflow, suggesting that diffusive broadening remained limited under the fabrication conditions used here. We next tested whether the embedded oligonucleotides remained chemically accessible and spatially resolved after polymerization. Hybridization of fluorophore-labeled complementary strands produced fluorescence localized to the intended domains (Figure 2D and 2E), demonstrating domain-resolved accessibility of the DNA anchors after continuous fabrication. Together, these results show that this platform can provide a scalable supply of uniform five-domain template microparticles whose DNA-defined domains remain spatially segregated and independently addressable for downstream post-fabrication encoding.

### Programmable Post-fabrication Encoding and Per-domain Coding Capacity

Having established in Figure 2 that the template particles retain domain-resolved DNA accessibility after continuous fabrication, we next asked whether these domains could be used to assign defined optical identities on a common particle template. To test this, a single batch of template particles was divided into four subsets and incubated with predefined cocktails of complementary oligonucleotides conjugated to the corresponding fluorophores: Alexa Fluor 405 (AF405), AF488, and AF647. This post-fabrication labeling produced four distinct multidomain fluorescence patterns (Figure 3A). This result shows that predefined barcode batches can be generated on demand from the same fabricated particles solely by changing the labeling mixture, thereby decoupling particle fabrication from barcode assignment.

**Figure 3.**
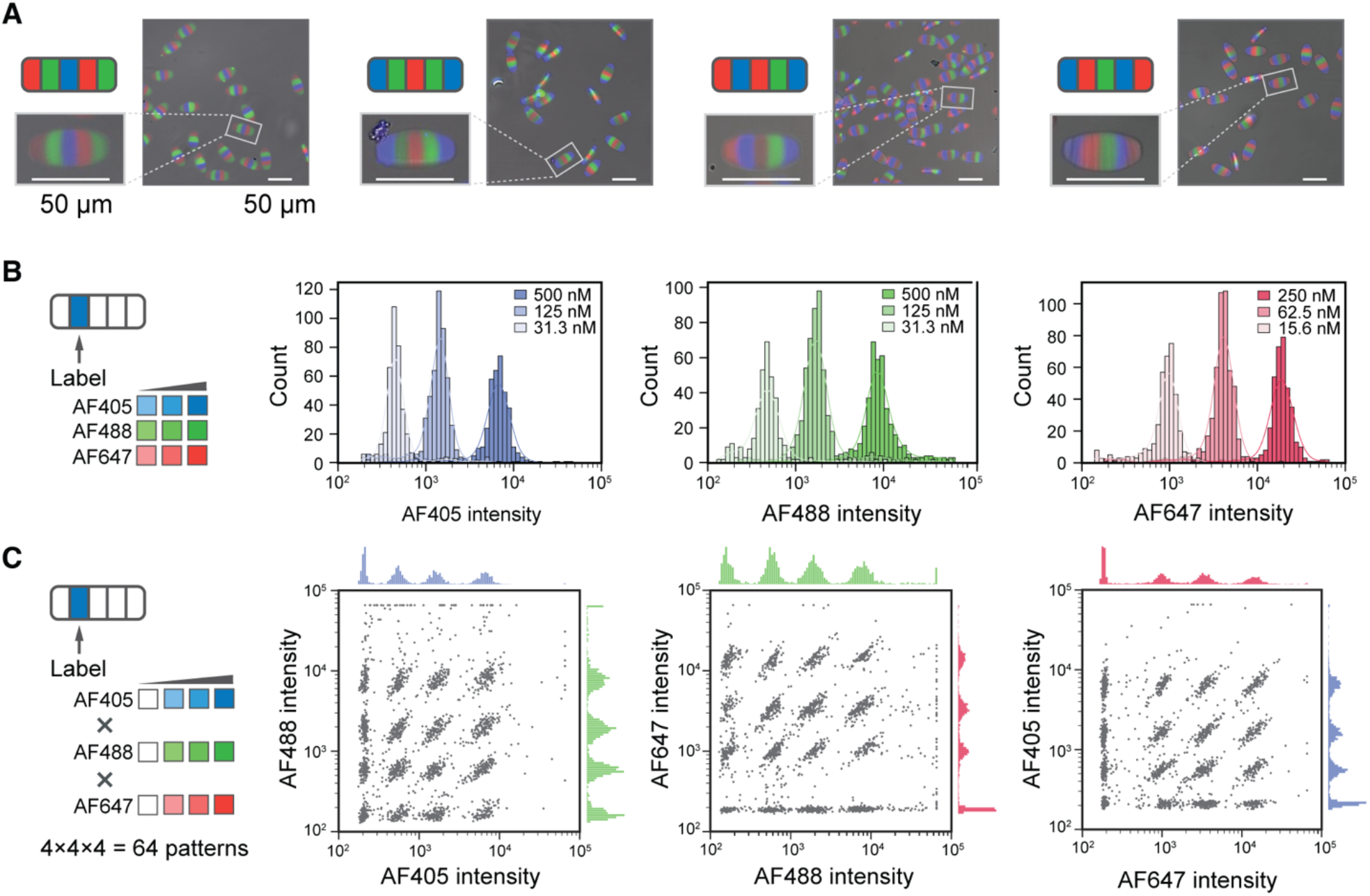
Programmable post-fabrication encoding by domain-selective hybridization. (A) Representative fluorescence overlays of distinct bead batches programmed to display predefined codes (blue: AF405, green: AF488, red: AF647). (B) Fluorescence intensity histograms showing three resolvable non-zero levels per fluorophore obtained by titrating staining concentrations (values indicated), enabling four addressable levels including the unstained state. (C) Pairwise intensity scatter plots for pooled 64 single-domain states (4^3^ combinations of three fluorophores, each with four addressable levels), showing grid-like cluster structure in AF405–AF488, AF488–AF647, and AF647–AF405 projections.

We then asked how many reliably distinguishable optical states could be assigned to an individual domain under standard fluorescence imaging. To estimate this per-domain coding capacity, we tuned the concentrations of AF405-, AF488-, and AF647-conjugated complementary oligonucleotides while maintaining the total oligonucleotide concentration constant. This produced three-resolvable non-zero intensity levels for each fluorophore (Figure 3B). Together with the unlabeled state, this encoding scheme defines four addressable levels per fluorophore, and their combination yields a total of 4^3^ = 64 possible three-color states per domain. When particles spanning this set were pooled, pairwise intensity plots showed the expected grid-like cluster structure for each of the 64 fluorophore combinations (Figure 3C). Extrapolating the experimentally resolved 64 candidate states per domain across five independently addressable domains gives a theoretical upper estimate of 64^5^ ≈ 1.1× 10^9^ barcodes. This value is an experimentally informed upper limit, whereas the operational library used below was restricted to a more conservative nine-state-per-domain codebook to improve decoding robustness.

### Decoding of a 59,049-code Library

We next asked whether the user-defined encoding scheme remains reliably decodable at large library size. To test this, we processed template particles by a split-pool workflow (Figure S1) with nine predefined per-domain spectral states to generate a library with 95 = 59,049 possible barcodes. We implemented the nine spectral states with nine distinct AF488 and AF647 labeling cocktails. AF405 was used not as a coding channel, but as an internal spatial fiducial: a stronger signal in domain 2 and a weaker signal in domain 4 provided a reference for domain ordering and segmentation. For decoding, individual particles were first identified in bright-field images, matched to the corresponding fluorescence images, and subjected to quality filtering (see Methods). The five domains were then segmented based on the AF405 fiducial signals, and the AF488 and AF647 intensities of each domain were extracted for 16,174 beads (Figure 4A).

**Figure 4.**
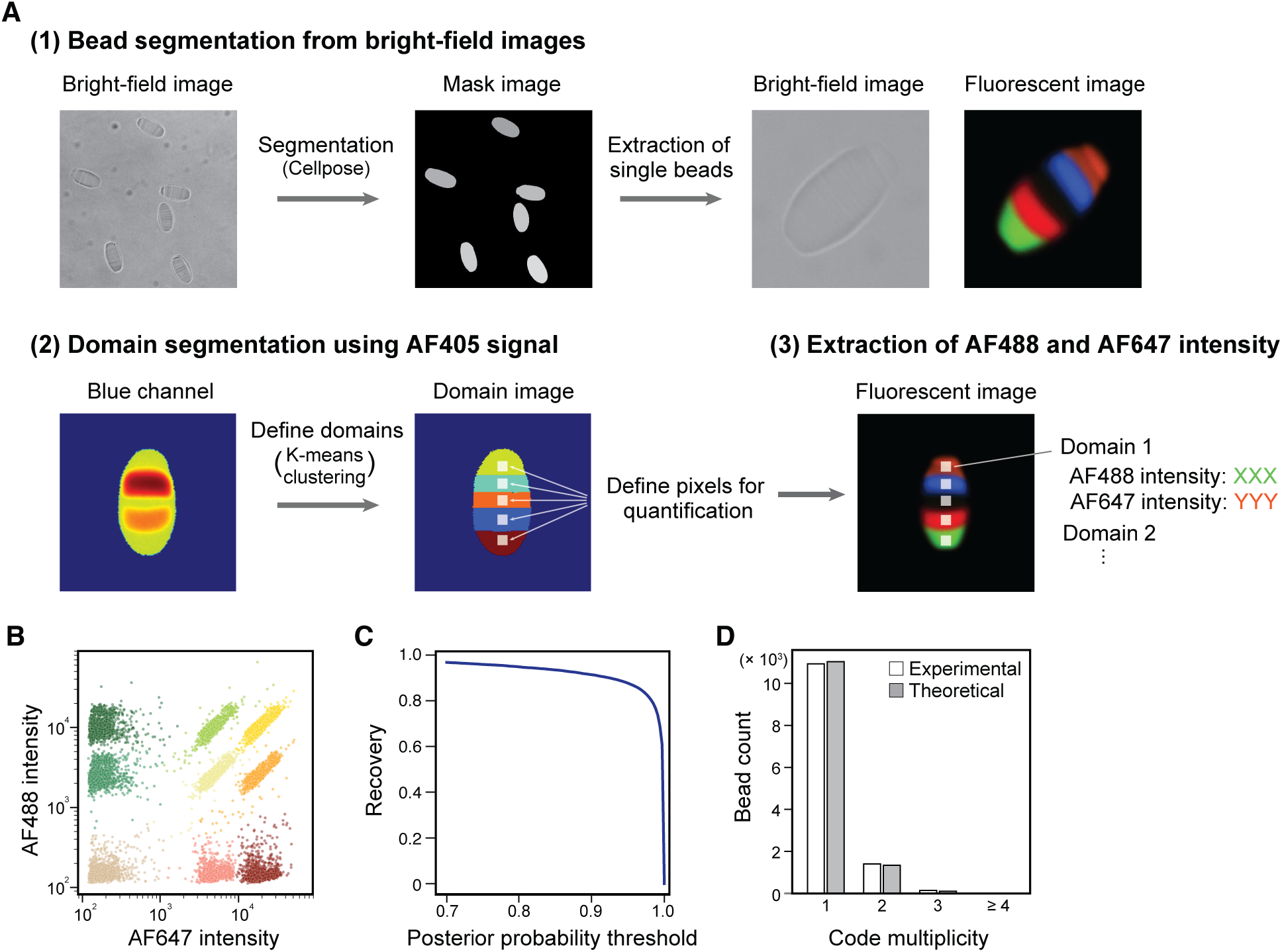
Decoding of a predefined 59,049-code library generated by split-pool labeling. (A) Image-processing pipeline for domain-resolved spectral intensity extraction: particle segmentation from bright-field images, domain ordering and segmentation using AF405 fiducial domains, and quantification of AF488/AF647 intensities from each of the five domains. (B) Representative AF647–AF488 intensity scatter plot for a single domain, showing nine spectral clusters assigned by a Gaussian mixture model (GMM). (C) Decoding recovery as a function of the posterior assignment probability threshold, with full-particle calls retained only when all five domains exceed the threshold. (D) Distribution of observed barcode multiplicity, defined as the number of beads assigned to the same decoded code, compared with the Poisson expectation for uniform random sampling from a 59,049-code codebook.

For each domain, we modeled the AF488 and AF647 intensity distribution using a Gaussian mixture model (GMM) with nine components corresponding to the nine spectral states (Figure 4B and S2). We then computed the posterior probabilities for each state assignment and used the maximum posterior probability as a measure of assignment confidence. Full-particle calls were retained only when all five domains exceeded a posterior threshold of 0.95. Under this stringent criterion, 14,198 out of 16,174 beads were successfully decoded, corresponding to an 88% recovery (Figure 4C).

To examine whether the split-pool process generated the 59,049-code library without strong representation bias, we examined barcode multiplicity, defined as the number of beads assigned to the same decoded code. The observed multiplicity distribution closely followed the Poisson expectation for uniform random sampling from a 59,049-member library (Figure 4D). This agreement suggests that the sampled population was broadly consistent with approximately uniform sampling from the predefined codebook without evident strong representation bias.

### Longitudinal Re-identification after 11 Days of Storage

For pooled screening applications, barcode readout must remain stable enough to support re-identification after long imaging intervals. We therefore re-imaged the library after 11 days of storage and constructed a positional ground-truth set of 10,654 matched bead pairs by identifying beads that remained at the same field positions in both imaging sessions. We then performed probabilistic matching using the full code-assignment probability vectors obtained from the GMM/Bayesian inference framework, allowing a match-confidence score to be computed for each candidate pair. When the pairwise match-confidence matrix was ordered according to the positional ground truth, the resulting heatmap showed a strong diagonal correspondence (Figure 5A and Figure S3), indicating that barcode readout remained sufficiently stable to discriminate correct bead identities after storage. Varying the match-certainty threshold allowed us to quantify the trade-off between re-identification accuracy and recovery (Figure 5B). At thresholds that maintained accuracy above 0.95, more than 7,800 beads were correctly re-identified, corresponding to over 73% of a positional ground-truth set of 10,654 matched bead-pairs. These results show that the DNA-linked spectral barcode readout remains sufficiently stable to support high-confidence longitudinal re-identification over at least 11 days, while ambiguous cases can be excluded by probabilistic confidence filtering.

**Figure 5.**
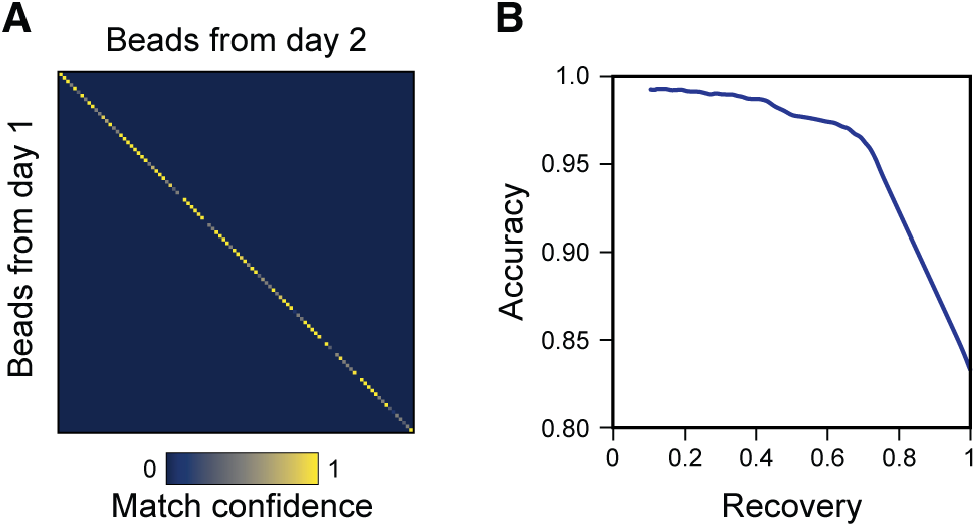
Longitudinal re-identification after 11 days of storage. (A) Heatmap of pairwise match-confidence matrix between the initial and 11-day datasets, shown for a 100-pair subset of the positional ground-truth set (10,654 matched bead pairs) and ordered according to the true bead correspondence. The strong diagonal indicates correct re-identification. (B) Accuracy–recovery curve obtained by thresholding the match-confidence score, wherein accuracy is defined as the fraction of retained matches consistent with the positional ground truth, and recovery as the fraction of the 10,654 matched bead pairs retained above threshold.

## Conclusion

We developed a barcoding architecture in which five-domain DNA-addressable hydrogel template microparticles are fabricated continuously at high throughput as a common template, and optical identities are assigned afterward by domain-selective hybridization. By separating template fabrication from barcode assignment, this platform shifts barcode diversification away from per-code reconfiguration of particle fabrication and into post-fabrication labeling and decoding. This design thus enables the generation of predefined batches of chosen codes from a single fabricated template and supports large, user-defined codebooks within a single-particle optical format.

Using contact-free multilaminar flow lithography with all-around three-dimensional sheathing, we continuously fabricated template particles at >10^6^ particles/h while preserving domain-level DNA addressability. On this scaffold, we demonstrated programmable post-fabrication encoding, established an experimentally informed upper-bound capacity of 64^5^ ≈ 1.1 × 10^9^ possible barcodes based on the resolved single-domain state space, and implemented a conservative 59,049-code (9^5^) library optimized for robust decoding. At a stringent posterior threshold, 88% of analyzed beads were successfully decoded, and more than 7,800 beads were correctly re-identified after 11 days at >0.95 accuracy.

Decoupling barcode assignment from fabrication does not remove manufacturing demands; it relocates them. Library performance still depends on uniform particle geometry, preserved domain segregation, accessible chemical addresses, and low defect rates. Relative to conventional deterministic optical barcodes, this architecture avoids code-by-code fabrication while retaining designed single-particle identities. Relative to stochastic combinatorial barcodes, it enables chosen barcode batches to be generated from a common scaffold rather than relying on which combinations happen to form. More broadly, it also differs fundamentally from sequencing-based strategies such as DNA-encoded libraries, in which identity is recovered by destructive sequencing. Here, DNA serves instead as a programmable chemical address for optical signals, enabling direct, in situ readout and longitudinal traceability.

Finally, the present implementation should be viewed as a modular foundation rather than an endpoint specific to DNA hybridization. Although barcode assignment here was performed by complementary oligonucleotide labeling, the same decoupled template architecture should in principle accommodate alternative post-fabrication chemistries, including covalent strategies compatible with harsh reaction conditions. More generally, the ordered address domains provide a natural basis for future coupling of barcode identity to functional payloads, offering a route toward user-defined code–payload mappings in pooled screening, phenotypic selection, and spatially resolved assay formats.

## Methods

### Materials

All oligonucleotide reagents (see also Table S1) were purchased from Integrated DNA Technologies (IDT) or Eurofins. Poly(ethylene glycol) diacrylate (Mn = 575; PEGDA575) was obtained from Sigma-Aldrich. Poly(ethylene glycol) (PEG200) was purchased from FUJIFILM Wako Pure Chemical Corporation. 2-Hydroxy-2-methylpropiophenone (HMPP) was obtained from Tokyo Chemical Industry (TCI). A blocking reagent CELLOTION was purchased from Takara Bio Inc.

### Microfluidic device fabrication

The microfluidic devices were fabricated in polydimethylsiloxane (PDMS; SILPOT 184, Dow Corning) using standard soft lithography. Two device layers were fabricated separately and manually aligned before bonding. The device has five sample inlets for the PEGDA solution, three sheath inlets, and one outlet. The five input samples join together first, followed by sequential confluence with bottom, top, and side sheath solutions. All the three sheath streams were split into two branches immediately after inlet entry, and merged with the sample stream from both sides.

### Theoretical Estimation of Inter-Domain Diffusion

To assess the potential for diffusive mixing between adjacent laminar flows prior to polymerization, we estimated the diffusion length (*L*_*d*_) using the Einstein-Smoluchowski relation: 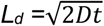, where *D* is the diffusion coefficient and *t* is the residence time. Based on the experimental flow rate (*Q* = 1.0 µL/min) and the cross-sectional area of the focused sample stream (*A* ~ 50 × 20 µm^2^), the flow velocity *v* is calculated as:

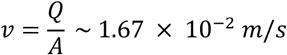

The distance from the confluence point to the UV irradiation zone is *d* = 700 µm. Thus, the residence time *t* is:

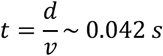

Using the diffusion coefficient of a typical short oligonucleotide (D ≈ 1.52× 10^-10^ m^2^/s), the diffusion length is calculated as:

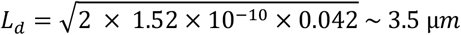

*Note*: We used the diffusion coefficient of DNA in a standard aqueous buffer. Since the viscosity of the 40% PEGDA solution is higher than that of the buffer, the actual diffusion coefficient in the microfluidic channel is likely lower, making this a conservative estimate of the broadening.

### Particle generation

All the solutions were sent to the device through PEEK tubes. The 40% PEGDA575 solutions with 5% HMPP and specific capture oligonucleotides (see Table S1) in DPBS were introduced to the device using a FLPG Plus (Fluigent, FLPG005J) and a LineUp Flow EZ (Fluigent, LU-FEZ-2000) at 6–15 mbar. To achieve contact-free 3D focusing, sheath solutions, 20% PEG200 in DPBS, are sent from syringes to the device using syringe pumps (Harvard Apparatus, PUMP 11 ELITE). The flow rates are 0.8–1.2 µL/min for the bottom and top sheath, and 2.4–3.0 µL/min for the side sheath. The laser beam (375 nm) to induce polymerization of the PEGDA solution was shaped into a rectangular profile of approximately 150 µm × 10 µm using a cylindrical lens and a spatial filter, and then tightly focused onto the channel by a high-NA (0.8) objective lens, enabling pulsed UV irradiation at 100–1000 Hz synchronized with the sample flow. The particles were washed with and kept in the TET buffer (20 mM Tris-HCl (pH 8.0), 0.5 mM EDTA, and 0.01% Tween 20) or TET-SC buffer (20 mM Tris-HCl (pH 8.0), 50 mM NaCl, 0.5 mM EDTA, 0.01% Tween 20, and 50% Cellotion).

### Domain-specific particle coloring (Figure 2)

The beads were incubated for 3 h in TET-SC buffer with 2 µM AF647-conjugated anti-DNA-1 and 2 µM of either AF488-conjugated anti-DNA-2, anti-DNA-3, anti-DNA-4, or anti-DNA-5. The incubation was performed at ambient temperature in the dark. After incubation, the beads were washed three times with TET-SC buffer (centrifuge: 100 × g, 3 min) and resuspended in the same buffer before imaging. The images were acquired using an INCell Analyzer 6000.

### Concentration-tuned encoding (Figure 3)

The beads were incubated overnight in TET buffer with a 1.5 µM oligonucleotide cocktail consisting of AF647-conjugated (0, 15.6, 62.5, or 250 nM), AF488-conjugated (0, 31.3, 125, or 500 nM), AF405-conjugated (0, 31.3, 125, or 500 nM), and non-labeled anti-DNA-2 oligonucleotides. Non-labeled oligonucleotides were added so that the total oligonucleotide concentration remained constant across all staining conditions. The incubation was performed at ambient temperature in the dark. After incubation, the beads were washed three times with TET buffer (centrifuge: 100 × g, 3 min), pooled, and resuspended in the same buffer before imaging. The images were acquired using an INCell Analyzer 6000 with autofocus based on the blue channel detecting the AF405 signal.

### Split-pool combinatorial encoding (Figure 4 and 5)

A library of encoded beads was generated via a split-pool method targeting 9 distinguishable states per domain. Initially, 1.8 × 10^5^ beads were divided into nine wells and incubated for 3 h with an oligonucleotide cocktail containing distinct ratios of AF647-, AF488-, AF405-, and non-labeled oligonucleotides in TET-SC buffer. The non-labeled oligonucleotides were supplemented to maintain a constant total oligonucleotide concentration across all staining conditions. After incubation, the beads were washed three times with TET-SC buffer (centrifuge: 100 × g, 3 min), pooled into a single reservoir, and re-aliquoted into nine wells for the subsequent layer encoding. This cycle was repeated for all designated domains, and details of the staining conditions are provided in Table S2.

The images were acquired using an INCell Analyzer 6000 with autofocus based on the blue channel detecting the AF405 signal. We correct chromatic aberration between the three channels with standard white agarose beads by applying an affine transformation using pyStackReg (v0.2.8), aligning the AF405 and AF647 channels to the AF488 channel. To exclude defocused images, we developed a three-layer CNN classifier trained on 52 and validated on 14 human-annotated bright-field images (35 focused and 31 defocused in total). Each input was a 256 × 256 pixel, intensity-normalized patch cropped from 2,048 × 2,048 pixel images divided into 8 × 8 patches. For each acquired image, the classifier predicted focus labels for all 64 patches, and the overall focus state was determined by majority voting. We used 628 images that were classified as in focus of the 800 images acquired.

### Bead and domain segmentation for decoding color-code (Figure 4 and 5)

The following image analyses were performed in Python 3.13. The analysis environment and all scripts are available at https://github.com/solabtokyo-org/PEGDABar-analysis.

Beads were segmented from focused bright-field images using Cellpose (v4.0.6)^[32]^ with the following parameters: flow_threshold=0.4, cellprob_threshold=0.0, tile_norm_blocksize=0, and batch_size=32. For each segmented bead, a bounding box was defined from the mask coordinates and expanded by 48 pixels on each side. The cropped image was then aligned by ellipse fitting and rotation so that the major axis became vertical.

Domain segmentation within each bead was performed using its bead mask and AF405 channel image. AF405-labeled domains—candidates for domains 2 and 4—were segmented by applying *k*-means clustering (*n* = 6) to the logarithmically transformed AF405 channel image. Morphological features (area, number of holes, circularity) were calculated for each cluster, and those with area ≥ 100 pixels, circularity of 0.3–0.9, and no internal holes were selected as AF405-labeled domains. Each selected domain was horizontally expanded to cover the full bead width based on its vertical coordinate range, yielding five domains per bead. For the remaining three domains, pixels with AF405 intensities greater than the mean + 15 σ of the background—calculated from regions outside the beads mask—were excluded from these candidate domains.

### Colorcode assignment (Figure 4 and 5)

For each domain of each bead, the integrated intensities of the AF405, AF488, and AF647 channels were calculated within a 5-pixel window centered at the domain centroid. Domain order was determined according to AF405 intensity (domain 2 > domain 4 > others). Outlier beads were excluded based on histogram-derived intensity gates: beads with domain 2 AF405 intensity between 4,175 and 26,053 or domain 4 intensity between 800 and 2,000 were retained.

We then employed a Gaussian mixture model (GMM) to assign AF488/AF647-defined color codes. The parameters of the GMM, with the number of expected color codes set as the number of components, were estimated from log_10_-transformed AF488 and AF647 intensities using the expectation–maximization algorithm as implemented in scikit-learn (v1.7.1). The fitted GMM defines, for each color code *c*, the likelihood p(x_*i,l*_|*c*) of observing the intensity vector x_*i,l*_ for bead *i* and domain *l*. Using the mixture weights as priors, we computed the posterior probability that domain *l* of bead *i* belongs to code *c* denoted as D_1_[*i,l,c*].

For each bead *i* and domain *l*, the color code with the highest posterior probability was assigned if this value exceeded a predefined threshold; otherwise, the bead was excluded as an outlier. Consequently, each bead was represented by a set of five domain-level color codes with associated posterior probabilities, forming its barcode identity.

### Bead matching (Figure 5)

To identify corresponding beads between images independently acquired on D1 and D2, we applied a matching algorithm based on joint-color-code assignment likelihood between every pair of bead *i* from D1 and bead *j* from D2. For each acquisition n, the posterior probability that the *k*-th domain of bead *i* has color code *c* was denoted as P_*nik*_(*c*). The matching likelihood between bead *i* and bead j was defined in the log domain as P(*i,j*) = Σ*l* log(D1[*i,l*,:] · D2[*j,l*,:]^T), representing the marginal log-likelihood of domain-wise color-code consistency between the two beads.

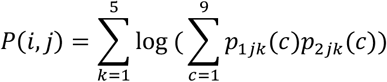

For each bead *i*, the values P(*i,j*) were normalized across all *j* using the softmax function so that ∑*j*P(*i,j*)=1 and the resulting softmax-normalized values were used as matching certainties.

### AI Usage

The authors used ChatGPT Pro (OpenAI) for English language editing. All content was reviewed and verified by the authors.

## Supporting Information

The authors have cited additional references within the Supporting Information.^[32]^

## Acknowledgements

This work was supported by JSPS KAKENHI grant numbers 25H01359 (to S.O.), 24K23030 (to A. E.), and 25K17926 (to A. E.); JST CREST grant number JPMJCR19H1 (to S.O.) and JPMJCR23B6 (to S.O.), JST GTex grant number JPMJGX23B1 (to S.O.), JST ASPIRE grant number JPMJAP2416 (to S.O.), and JST ACT-X grant number JPMJAX2534 (to A.E.); The Uehara Memorial Foundation (to S.O.); UTEC-UTokyo FSI Research Grant Program (to S.O.); Takeda Science Foundation (to S.O.); and Nakatani Foundation (to S.O.).

## Supplementary Figures

**Figure S1.**
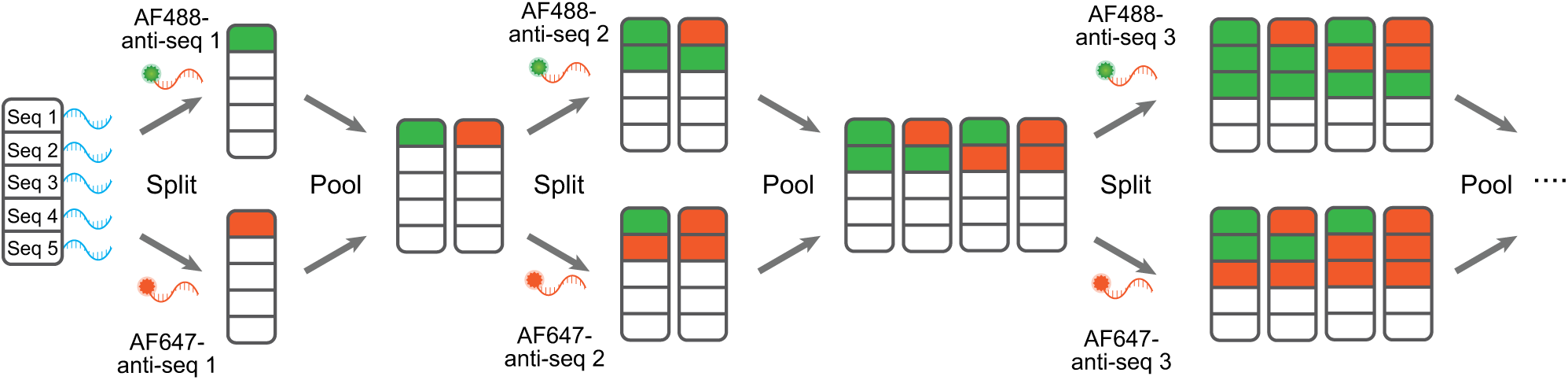
Schematic illustration of split pool-based particle coloring.

**Figure S2.**
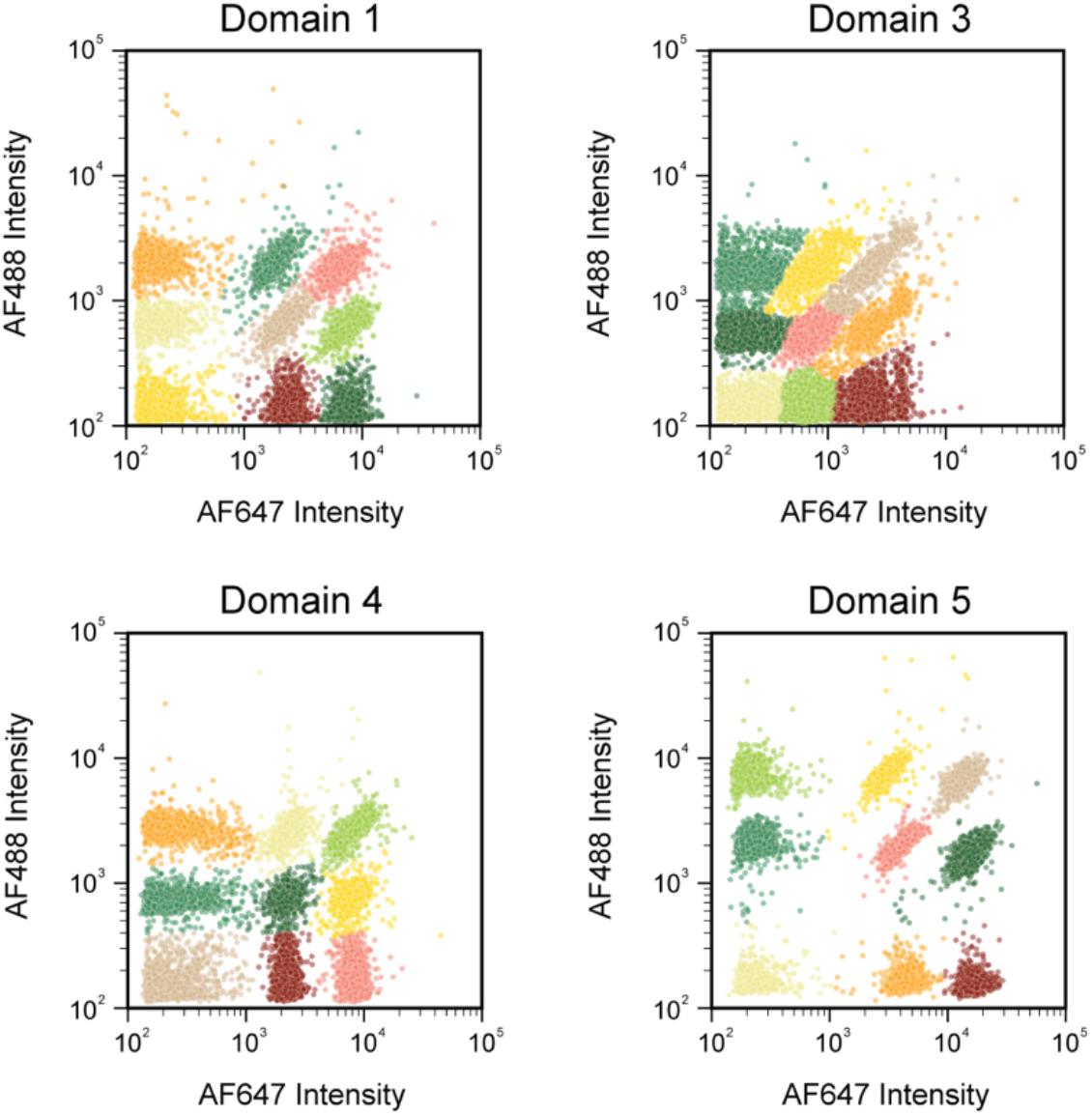
2D intensity scatter plot of AF488 and AF647 signals from the domains 1, 3, 4, and 5 of the beads, exhibiting nine distinct clusters. The color represents the nine clusters assigned by GMM.

**Figure S3.**
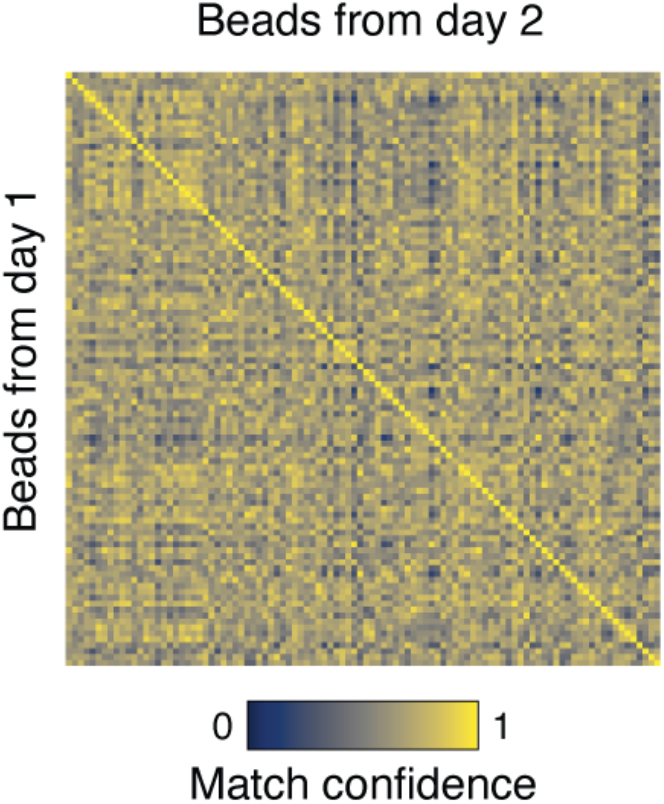
Heatmap of matching certainty scores before softmax normalization between beads imaged on Day 1 and Day 2, highlighting 100 beads out of 10,654 beads.

## Supplementary Tables

**Table S1.**
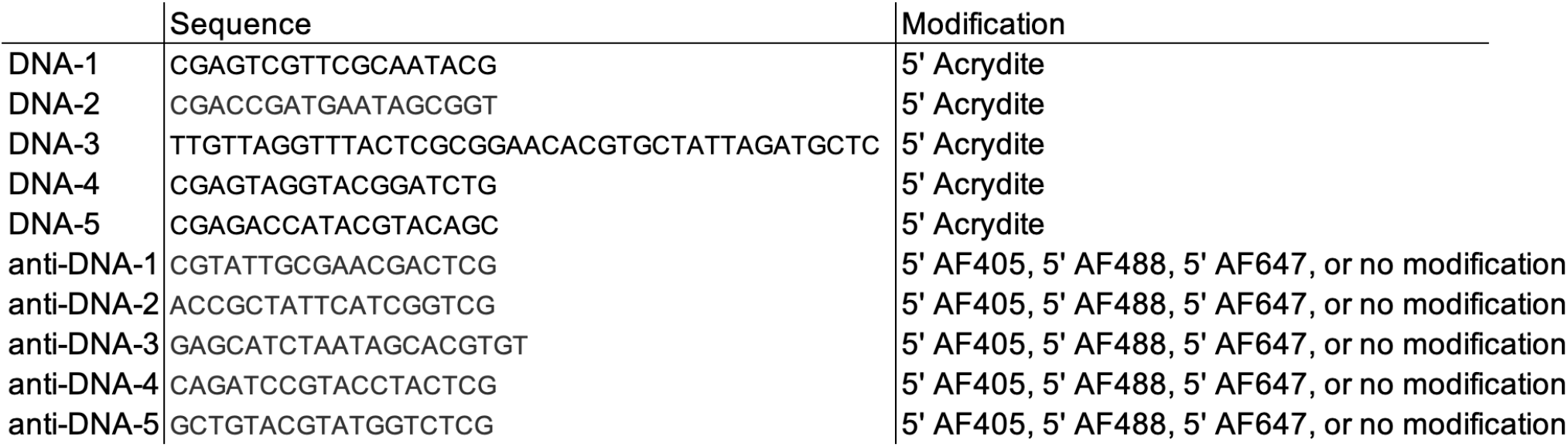
List of the sequence of the oligonucleotides used in this paper.

**Table S2.** Staining conditions used for split-pool library generation.

| Anti-DNA-3 (round 1) |  |  |  |  | Anti-DNA-4 (round4) |  |  |  |  | Anti-DNA-2 (round5) |  |  |  |  |
| --- | --- | --- | --- | --- | --- | --- | --- | --- | --- | --- | --- | --- | --- | --- |
| Anti-DNA-5 (round 2) |  |  |  |  |  |  |  |  |  |  |  |  |  |  |
| Anti-DNA-1 (round 3) |  |  |  |  |  |  |  |  |  |  |  |  |  |  |
|  | AF405 | AF488 | AF647 | Non-label |  | AF405 | AF488 | AF647 | Non-label |  | AF405 | AF488 | AF647 | Non-label |
| 1 | 0 | 0 | 0 | 3000 | 1 | 250 | 0 | 0 | 2750 | 1 | 500 | 0 | 0 | 2500 |
| 2 | 0 | 0 | 125 | 2875 | 2 | 250 | 0 | 125 | 2625 | 2 | 500 | 0 | 62.5 | 2437.5 |
| 3 | 0 | 0 | 500 | 2500 | 3 | 250 | 0 | 500 | 2250 | 3 | 500 | 0 | 250 | 2250 |
| 4 | 0 | 250 | 0 | 2750 | 4 | 250 | 250 | 0 | 2500 | 4 | 500 | 125 | 0 | 2375 |
| 5 | 0 | 250 | 125 | 2625 | 5 | 250 | 250 | 125 | 2375 | 5 | 500 | 125 | 62.5 | 2312.5 |
| 6 | 0 | 250 | 500 | 2250 | 6 | 250 | 250 | 500 | 2000 | 6 | 500 | 125 | 250 | 2125 |
| 7 | 0 | 1000 | 0 | 2000 | 7 | 250 | 1000 | 0 | 1750 | 7 | 500 | 500 | 0 | 2000 |
| 8 | 0 | 1000 | 125 | 1875 | 8 | 250 | 1000 | 125 | 1625 | 8 | 500 | 500 | 62.5 | 1937.5 |
| 9 | 0 | 1000 | 500 | 1500 | 9 | 250 | 1000 | 500 | 1250 | 9 | 500 | 500 | 250 | 1750 |
|  |  |  |  | (µL) |  |  |  |  | (µL) |  |  |  |  | (µL) |

